# Optical Clearing with Tartrazine Enables Deep Transscleral Imaging with Optical Coherence Tomography

**DOI:** 10.1101/2024.10.25.620343

**Authors:** Amit Narawane, Robert Trout, Christian Viehland, Anthony N. Kuo, Lejla Vajzovic, Al-Hafeez Dhalla, Cynthia A. Toth

## Abstract

Suprachoroidal injections are a relatively new method of drug delivery to treat retinal disease. At present, it is difficult to visualize the distribution of injection-delivered product beneath the sclera into the suprachoroidal space. Imaging the suprachoroidal space with OCT is hindered by scattering of light from densely packed collagen fibers of the sclera, limiting depth penetration of the OCT light. In this Letter, we demonstrate the first use of tartrazine (Yellow 5) as an optical clearing agent for suprachoroidal research imaging with OCT in an *ex vivo* eye model. Our results show that this clearing agent dramatically improves visualization of the choroid and suprachoroidal space with transscleral OCT and allows for improved imaging of the location and extent of delivered suprachoroidal fluid. We believe these methods will enable use of optical techniques such as OCT to image through a variety of previously optically inaccessible, highly scattering tissue samples.

---

Retinal diseases such as age-related macular degeneration, diabetic retinopathy, and uveitic macular edema are leading causes of visual impairment, with rising global prevalence^1–3^. Intravitreal injections of anti-VEGF agents and/or steroids have been effective in controlling vascular leakage and often restoring visual acuity, but they require repeated injections. Novel treatments such as intravitreal reservoirs or subretinal injections of gene therapies can be effective over a longer period, but may cause glaucoma and cataract (in the case of steroids) or incur the cost and complexity of delivery in a microsurgical setting (in the case of subretinal gene therapy)^4^. This has led to development of ocular therapeutics that can be delivered by transscleral injection into the suprachoroidal space in an office-based procedure. Triamcinolone suspension for injection into the suprachoroidal space was approved in 2021 by the U.S. Food and Drug Administration for treatment of uveitic macular edema, and clinical trials are underway using this delivery route for other posterior segment diseases. A limitation of this procedure is that clinicians are not able to obtain real-time feedback on the location and spread of the delivered product; rather, postmortem animal studies, or heat map imaging in the case of human delivery, are used to estimate this^5,6^.

Optical coherence tomography (OCT) has widespread use in ophthalmology and allows for micron-scale depth resolution of the anterior eye^7,8^. We have previously demonstrated the potential for intraoperative OCT systems to measure drug delivery volumes during subretinal injections^9^. We hypothesize that this technology also has the potential to visualize drug delivery to the suprachoroidal space in real-time. To do this, however, requires transscleral OCT imaging of the suprachoroidal space. This in turn requires ballistic photon travel through the sclera, a tissue composed of dense, randomly oriented collagen^10^. This medium is highly scattering at the near infrared (NIR) wavelengths utilized by OCT systems^11^, limiting the imaging depth to 1-2mm, which is insufficient to visualize the suprachoroidal space^12,13^.

To improve OCT signal return from greater depths in scattering tissue, several hardware-based approaches have been applied towards, including separated illumination and collection pathways to recover multiply scattered light^14^, and ultrasound-based mechanical clearing of tissue^15^. However, these methods require significant augmentation of the imaging system. As an alternative, chemical approaches involving biocompatible tissue clearing agents have been investigated for potential in OCT imaging, including various sugars, glycerol, propylene glycol, and dimethylsulfoxide (DMSO)^12,16–18^. Their mechanism consists of increasing the concentration of the clearing agent within the interstitial fluid of the tissue via topical exposure of the tissue to a solution of the agent. The accompanying increase in the refractive index of the interstitial fluid results in optical clearing and reduced scattering. However, the efficiency of the refractive index change for these agents is poor, requiring high concentrations within the tissue to achieve substantial optical clearing. Recently, tartrazine (Yellow 5) has been shown to have potential as a more efficient biocompatible optical clearing agent for biomedical tissue imaging with brightfield, fluorescence, laser speckle contrast, and second harmonic generation microscopy^19^. Motivated by these findings, we endeavored to investigate whether tartrazine could be used to improve OCT depth penetration in highly scattering tissue such as the sclera, and in this case, to facilitate improved visualization of suprachoroidal injection dynamics.

In this Letter, we describe transscleral imaging with and without tartrazine using a custom-designed handheld telecentric OCT probe designed for high resolution imaging of the anterior eye. We first evaluated our OCT probe for imaging the anterior segment by correlating imaging results from *ex vivo* human eyes to known anatomical structures seen on histology. Then, we demonstrated, in *ex vivo* porcine eyes, that tartrazine optically clears scleral tissue for our infrared imaging bandwidth to improve the depth penetration of OCT imaging through the sclera. To our knowledge, this work presents the first use of tartrazine as a non-invasive optical clearing agent for OCT. We believe these methods will enable OCT studies in thick, scattering samples that were previously optically limited or inaccessible.

To attempt transscleral imaging of the suprachoroidal space, we developed a purpose-built swept-source OCT system, including a telecentric sample arm with custom-designed optics and opto-mechanics (Fig. 1). The probe is capable of operating in both handheld and table-mounted modes and provides a 25 mm working distance, lateral resolution of 12.7 μm (Airy radius), and lateral field-of-view >18 mm. The OCT engine employs a swept-source laser operating at 1040 nm with a 104nm bandwidth and 200 kHz sweep rate (Excelitas, Axsun Technologies Inc), delivering an axial resolution of 5.92μm (in air) and average sample arm power of 1.23 mW. This OCT system is well-suited for use in imaging and evaluating suprachoroidal injections both *ex vivo* and in the clinic.

**Fig 1.**
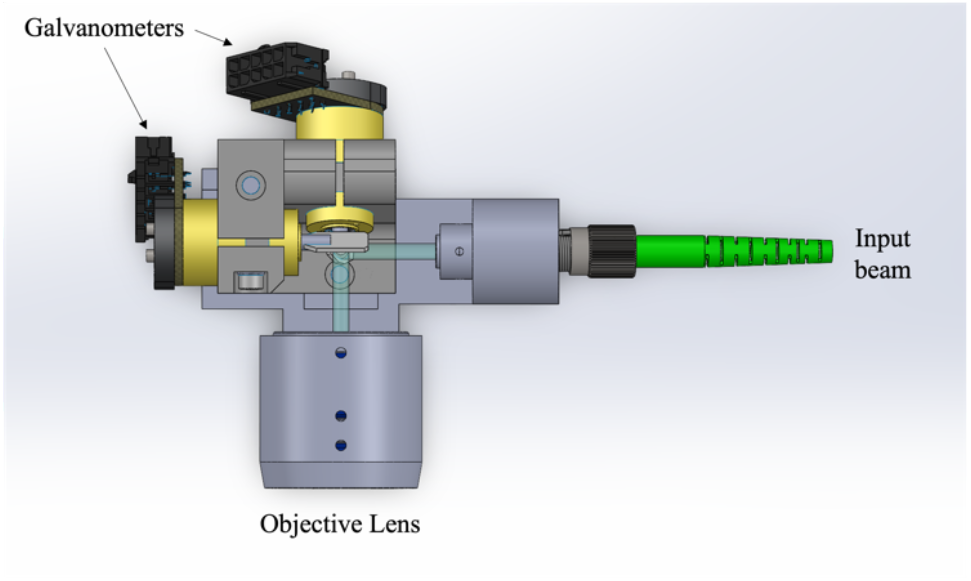
Schematic of custom handheld anterior segment probe, including galvanometers for scanning and objective with custom lenses optimized for telecentric beam focusing. Probe is contained within custom 3D printed casing designed with a handheld grip.

Imaging suprachoroidal injections with our OCT system necessarily involves imaging through the sclera at the correct region of interest in the anterior segment. It is therefore important to be able to demarcate different structures in the anterior segment within the OCT images to correctly identify the suprachoroidal space and evaluate placement of the injected agent. To this end, we performed a preliminary experiment on post-mortem donor human donor eyes. Eyes were marked with sutures that acted as fiducial markers between images. The eyes were imaged across the limbus in the superotemporal quadrant with our OCT system. Our scan protocol imaged a field of view (FOV) of 13 × 13 mm, with 1000 A-scans/B-scan and 128 B-scans/volume, and 8x repeated B-scans at each location for averaging. We then fixed the eyes in formalin for 24 hours followed by paraffin embedding, sectioning and hematoxylin and eosin (H&E) staining. The sutures were used to align B-scans with their corresponding histological sections. Figure 2 shows an example of correlations between an OCT B-scan and its corresponding H&E section using known anterior segment anatomical features ^20^. Note that while many structures can be readily identified on OCT, it is still difficult to resolve the boundaries of the suprachoroidal space and the choroid with conventional transscleral imaging.

**Fig 2.**
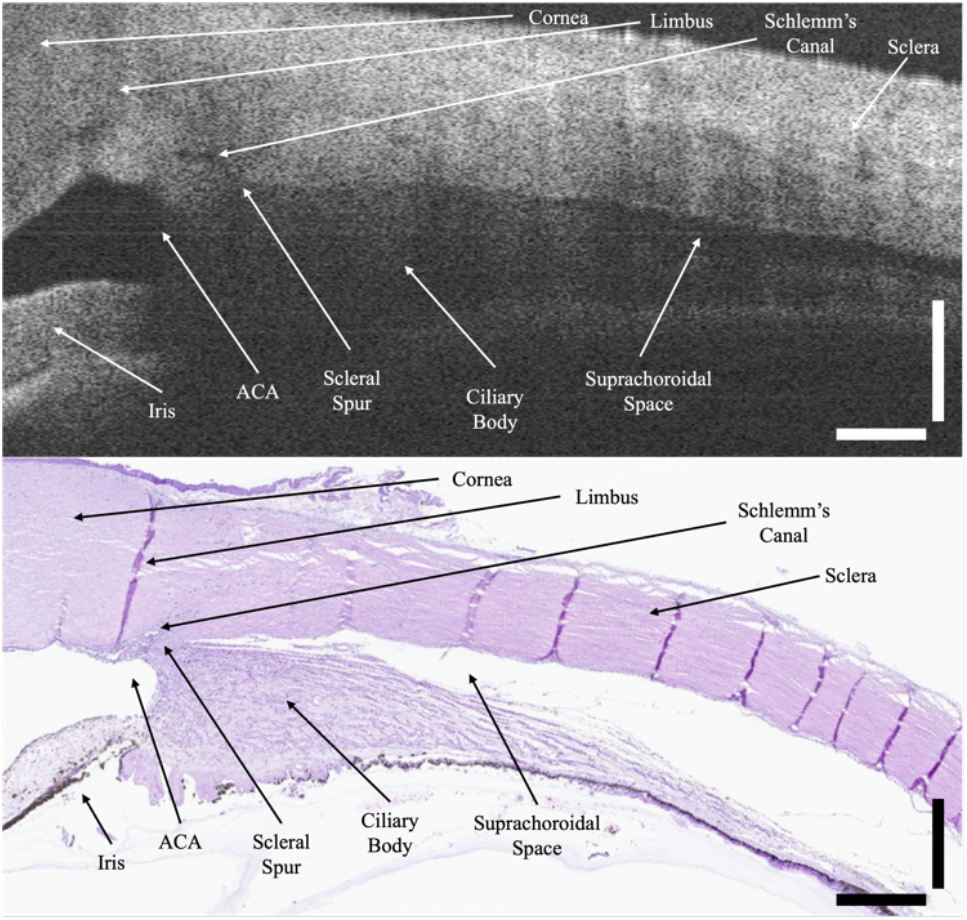
Correlation of anatomical features of the anterior segment between a B-scan from our OCT system and the corresponding H&E section. Note that the suprachoroidal space is visible likely due to choroidal separation in the donor eyes during processing and transit, and the conjunctiva and Tenon’s complex was removed prior to imaging. ACA: anterior chamber angle. Scale bar 0.5mm in tissue.

To test optical clearing in OCT, a pair of *ex vivo* porcine eyes, derived from the same animal to ensure comparability, were used in this experiment. Porcine eyes are preferred for iterative studies as they are readily available fresh from an abbatoir. There is also typically a delay post-mortem in obtaining donor human eyes, during which choroidal and retinal separation is common (as seen in Fig. 2).

This makes it difficult to determine whether the suprachoroidal space is visible due to injection or tissue separation. Porcine eyes can be acquired immediately after death, which makes them a superior model for studying real-time suprachoroidal injections. Although porcine sclera from small animals has been reported to be comparable in thickness to that of the human^21^, the sclera of our specimens have been thicker (consistent with the larger source pigs from 200-360kg), ranging from 1 mm thick at 8 mm posterior to the limbus in the superotemporal quadrant, to 2 to 3 mm thick across most other regions.

One globe was immersed in a 0.4 M tartrazine solution. At 0.4 M, 1.6 Osm, this tartrazine solution is hyperosmotic to tissue (1.6 Osm > 0.3 Osm)^22^. As a hyperosmotic solution, there is a dehydrating effect on the tissue that can confound results with increased imaging depth. To distinguish between the effect of dehydration and refractive index matching, our second eye pair served as a control, and was immersed in a saline solution of equal osmolarity to the tartrazine solution (0.8 M NaCl, 1.6 Osm). Each eye was immersed for a total of 120 minutes, with imaging timepoints at 0, 10, 20, 30, 60, and 120 minutes. At each timepoint, the immersion solution was temporarily removed for approximately 1 minute to expose the eye for imaging. Both eyes were imaged in the same temporal region.

Figure 3 shows a representative B-scan from each eye, one in saline and one in tartrazine, captured and processed with the same protocol as above at each time point but with an increased FOV of 18 × 18 mm. While the eye in saline showed minimal change from start to finish, the eye in tartrazine showed dramatically increased depth penetration of OCT light through the sclera over time. At the 2-hour mark, we achieved near full visualization of the choroid through the sclera, with individual choroidal vessels visible that were completely unseen in the control eye.

**Fig 3.**
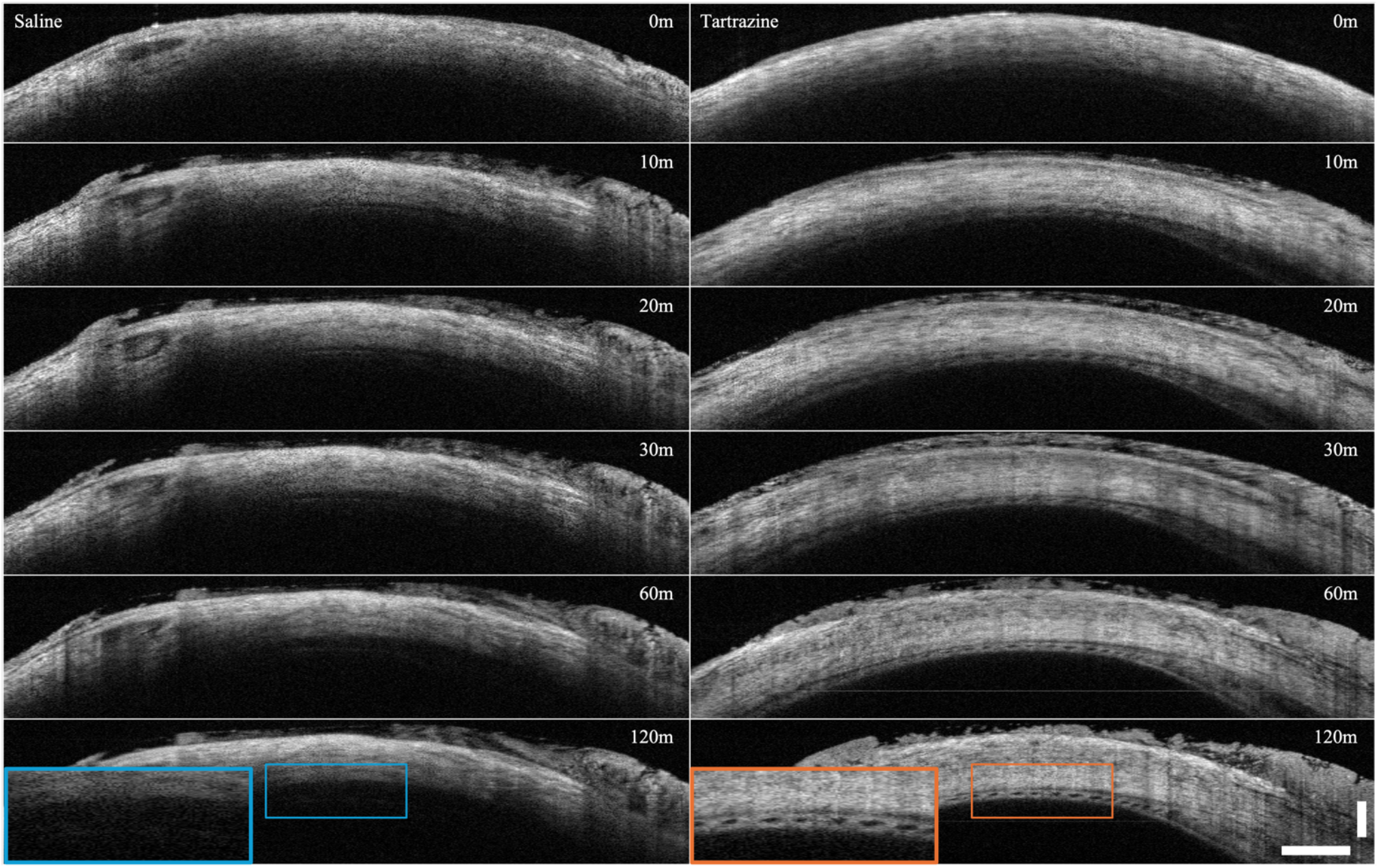
OCT imaging of one pair of *ex vivo* porcine eyes derived from the same animal, targeting the sclera on the temporal side for each. One eye from the pair (left column) is immersed in saline, the other (right column) in tartrazine. Each eye was imaged at the same location at 0, 10, 20, 30, 60, and 120 minutes. Blue and orange insets show revealed choroid boundaries and large choroidal vessels in the tartrazine eye. Scale bar 1mm in tissue.

To more directly compare the intensity changes over depth with optical clearing by tartrazine, we compared the intensity profile in depth within each image. The central 10 A-scans of each image in Figure 3 were averaged, and the smoothed result was cropped to extend from the hyperreflective scleral surface past the lower boundary of the sclera (Fig. 4). Here we see increased signal return at greater depths past the scleral surface in the tartrazine exposed eye than we do in the control eye. We note that the dip in intensity in the shallow parts of the sclera is expected; as the tissue is surrounded by material with a high refractive index, less light is reflected by the shallower parts of the sclera, and instead penetrates further into the tissue before being back-scattered at a deeper layer.

**Fig 4.**
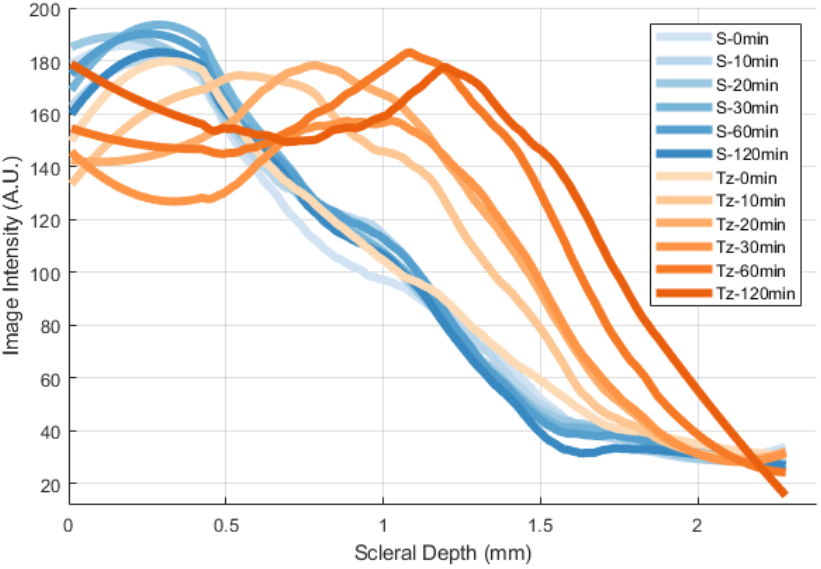
Plotted depth intensity profiles for each image depicted in Figure 3. Profiles derived as the central 10 A-scans of each image averaged and smoothed, then plotted from the hyperreflective scleral surface into the vitreous below. Legend: S – saline; Tz – tartrazine.

In order to determine the impact of optical clearing with tartrazine to image suprachoroidal drug delivery, we performed a suprachoroidal injection on porcine eyes with and without exposure to tartrazine. Injections were 0.1mL of saline or steroid solution and were performed using the SCS Microinjector (Clearside Biomedical). OCT volumes were acquired before and after each injection with the same protocol described above. Representative B-scans from the OCT volumes after injection are shown in Figure 5. While the lower boundary of the suprachoroidal space is difficult to visualize in the first eye, the same boundary is clearly seen once the sclera has been optically cleared with tartrazine.

**Fig 5.**
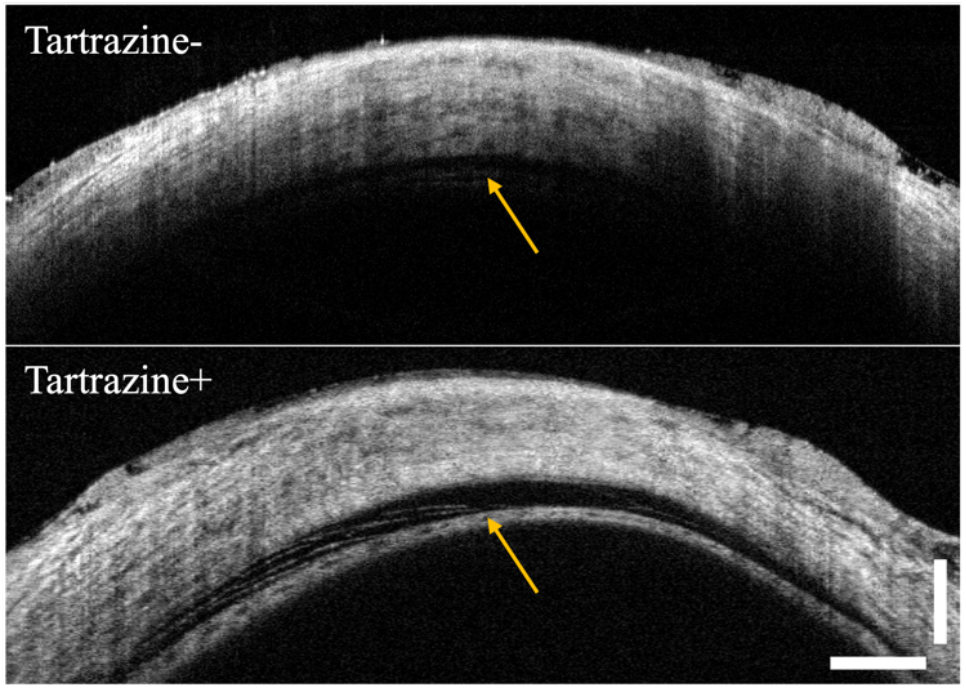
Representative B-scans from OCT volumes captured after injection into the suprachoroidal space in *ex vivo* porcine eyes without (top) tartrazine and with (bottom) 2-hour tartrazine immersion. Yellow arrows indicate location of choroidal boundary with the suprachoroidal space above it after injection, in contrast to the absence of the space prior to injection (Fig 3). The signal in the tartrazine eye reveals strands of reflective tissue bridging the suprachoroidal space (left of the arrow) and the broad extent of choroidal separation. Scale bar 1mm in tissue.

In this work, we have achieved the first demonstration of optical clearing with OCT using highly absorbing molecules. Our results show that the optical clearing methods described in^19^ are extendable to OCT imaging, and can likely be applied to any reflectance-based imaging modality. We have also demonstrated that our custom handheld anterior segment OCT system is capable of scleral imaging and visualization of the suprachoroidal space, and of the choroid and choroidal vasculature with optical clearing agents.

While tartrazine was demonstrated to have a large effect on optical clearing in scleral OCT imaging, it is possible that further improvement could be seen with the application of other food-grade dyes similar to tartrazine but with a more red-shifted absorption spectrum, such as Allura-Red (Red 40), which may result in greater optical clearing effect in the NIR region of OCT imaging.

Here we investigated *ex vivo* tissues, and the logical next step would be to conduct *in vivo* studies on animal models to validate the method in a living system. It is expected that there may be challenges with living vasculature removing the clearing agent more rapidly than in *ex vivo* samples, however with the potential benefit of improved visualization of the dynamics of living tissue and injection procedures. We anticipate continuation of this work will allow us to further study transscleral imaging of the suprachoroidal space, with the eventual goal of improving visibility of suprachoroidal injection therapies in humans.

## Funding

National Institutes of Health (R01 EY023039, U01 EY028079, R01 EY025009, F31 EY035168, Core Grant P30 EY005722), Research to Prevent Blindness (Unrestricted to Duke Eye Center).

## Acknowledgments

We would like to acknowledge Joseph A. Izatt for his invaluable support and mentorship in the earlier stages of this project. We would also like to thank Michelle McCall for administrative support.

## Disclosures

CV: Theia Imaging (I,P,E), ANK: Leica Microsystems (P), LV: Alcon Inc. (F,C), Clearside Biomedical (C), Novartis (R,C), AHD: Theia Imaging (I,P,E), Leica Microsystems (P) Horizon Surgical (I,P,E) CAT: Carl Zeiss Meditech (R), Theia Imaging (I,C), Emmes (C)

## Data Availability

Data underlying the results presented in this paper are available from the authors upon reasonable request.

